# Modelling the displacement and coexistence of clonal lineages of *Phytophthora infestans* through revisiting past outbreaks

**DOI:** 10.1101/2023.03.08.531797

**Authors:** Chih-Chiang Huang, Edward C.Y. Liew, Justin S. H. Wan

## Abstract

The continuous changes in the lineage proportions of populations in the clonal plant pathogen *Phytophthora infestans* on potato and tomato crops have been perplexing to researchers and disease managers. Sudden outbreaks of newly emergent genotypes are often associated with these rapid composition changes. Modelling can predict the persistence and displacement of pathogen genotypes with differential fitness among hosts. Building upon previous models, we combined analytical and simulation methods to model the outcome of interactions between competing lineages on multiple hosts. Model inputs include pathogenesis parameters, and the outputs are fitness and lineage proportions within each host. Analytical solutions yielding complete displacement, partial coexistence-displacement, and complete coexistence were described. In a retrospective study, the lesion growth rate and sporulation density of *P. infestans* lineages on potato and tomato from pathogenicity trials were used as inputs. Output lineage frequencies were compared with historical epidemiological situations to check model accuracy. The results showed that pathogenesis traits measured from empirical trials could simulate lineage constituents on potato and tomato, and estimate genotypic fitness with reasonable accuracy. The model also showed promise in predicting ongoing lineage displacements in the subsequent year or few years, even when the displaced lineage was still highly prevalent during the time of isolation. However, large uncertainties remain at temporal-spatial scales owing to complex meta-population dynamics in some regions and adaptation to local environmental factors. This simulation model provides a new tool for forecasting pathogen compositions, and can be used to identify potentially problematic genotypes based on pathogen life-history traits.

## Introduction

Many of the most economically significant plant pathogens are chiefly asexual in the mode of reproduction. Asexual reproduction in plant pathogens allows propagules to very rapidly infect available hosts in an area (e.g., in a field). Agricultural clonal plant pathogens have unique characteristics in their population dynamics and epidemiological properties. These characteristics include changes in the frequencies of clonal lineages with each season, or temporary stability in the dominance of certain lineage (Kohn 1995). A lineage is defined as a multilocus genotype grouping of clones (i.e., MLG), and frequency is the proportion of the MLG within a given pathogen population. The study of the variability in the MLG compositions within pathogen populations is important because of their link to the epidemiology of outbreaks. Some invasive MLGs that are able to spread rapidly through a landscape may be associated with severe disease, fungicide resistance or overcoming of host defences. Among these, the most destructive are the emergent lineages of invasive *Phytophthora* sp., including potato and tomato late blight caused by *P. infestans*, sudden oak death pathogen *P. ramorum*, and Panama disease caused by *Fusarium oxysporum* f. sp. *cubense* (‘Foc’, Ploetz 2005). Recurrent emergences in the globally distributed pathogens are linked to fluxes at the global or the regional (continental meta-population) level. Due to the massive scale of the problem, it is necessary to develop appropriate tools and management strategies.

*Phytophthora infestans* is a serious oomycete phytopathogen that has seen continual dramatic changes to the genotypic frequency of populations on potato and tomato hosts (Yoshida et al. 2013; Fry 2020). It is an almost completely asexual species and an obligate pathogen that cannot survive long outside its hosts. Dispersal is primarily aerial (especially with high humidity) and spores can be transmitted many kilometres (Fry and Grünwald 2010). Population growth in the pathogen is often explosive and outbreaks can happen extraordinarily fast (Fry and Grünwald 2010). While mathematical models of disease severity based on weather conditions, fungicide application and cultivar selection have been used to accurately predict disease (e.g., LATEBLIGHT for potato late blight; Andrade-Piedra et al. 2005), there are only a few models on the displacement and coexistence of competing genotypes in clonal plant pathogens. There are various models on the competition between two pathogen genotypes with fitness based on sporulation and infection parameters (e.g., in rust fungi, Newton et al. 1998). Models usually include mixed mathematical and simulation modelling (Mailleret et al. 2011). Yuen (2012) previously modelled the competition in *P. infestans* between two lineages (termed ‘population’ in the model), using a Lotka-Volterra model to simulate whether the displacement of a resident lineage by a new lineage occurs in a host species within a single season. However, both potato and tomato are major hosts of *P. infestans* worldwide. A model that deals with multiple lineages and hosts is necessary to investigate the interactions between genotypes of differing qualities. For example, generalist strains can infect multiple host species with equal aggressiveness but specialists may only infect a single host species. A model that can simulate genotype frequencies based on pathogenicity traits may help identify potentially invasive emergent strains.

In this paper, we develop a model that predicts the frequency of lineages within populations on multiple hosts. Fitness of each lineage on each host was mainly based on lesion growth rate and relative sporulation capacity, which are assumed to be inert and does not change with time. Lineages can be competing taxa (species) within closely-related groupings or MLGs within a species.

## Methods

### Model derivation

This model considers the co-existence or displacement of *P. infestans* lineages with varying pathogenicity on two hosts (e.g., potato and tomato). In this paper, complete displacement of a lineage is considered the case where that lineage is virtually absent in the population (e.g., <1% in the population). The main input variables are lesion growth rate (*LGR*) and relative sporulation capacity (*SD*) of each lineage on each host, which determines fitness (*F*) and cross-host transmission (*β*) for each lineage. Hosts are assumed to be ubiquitous agricultural crops so the effects of host density are not considered explicitly. Time is unscaled with discrete steps (iterations). Cross-host transmission follows the effects of competitive interactions (*c*) on fitness in a selection-transmission demographic model. Lesion area (*LGR*) is the key component in competitive interactions (e.g., Barrett et al. 2021). Genotypes are assumed to be completely clonal with no mutation so pathogenesis traits do not change with time. In each iteration *i*, the new pathogen population size (*N*) within each host is changed according to fitness adjusted by density dependence and competitive effects, *N*_(*i*+1)_ = *F·N_i_* This is followed by density dependent cross-host transmission (there is no within-host species transmission). We assume that fitness is also mildly density dependent, where the available space and resources are naturally diminished as the pathogen population size increases. This density dependent fitness yields an upper limit for *N*. Relative sporulation capacity (i.e., proportional) was used instead of absolute *SD* due to the measure of the unscaled *N*. For *k* lineages, *SD* for lineage *l* within host *h* is defined as: *SD_l,h_* = *absolute SD_l,h_ / ∑(absolute SD_k,h_*). Hence, the model output for *N* is deterministic in regards to the fitness as defined by *LGR* and *SD*. That is, all values for the starting population size (*N*_0_) result in the same final lineage proportion values (each lineage needed to be initially present on either host). Fitness (*F*) of the pathogen lineages on the two hosts is described in Montarry et al. (2010). For each lineage on each host (lineage *l* on host *h*):

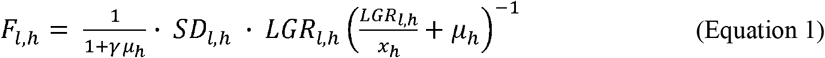

Where the constants *γ* is the latent period (days); *μ* is inverse time available for infection (inverse season length); *x_h_* is the carrying capacity on the tissues of host *h*, *SD_l,h_* is the relative sporulation capacity and *LGR_l,h_* is lesion growth rate of lineage *l* on host species *h*, respectively.

The form of density dependence follows Maynard Smith and Slatkin (1973) and Gomulkiewicz et al. (1999), which assumes that fitness decreases as the population size approaches some carrying capacity. The population size following density dependent regulation (*N’*) is defined by:

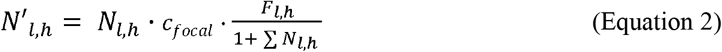

*N* is the original population size of pathogen lineage *l* on host *h*. The effect of competition between lineages occurs within hosts, and is modelled using relative *LGR* (i.e., the space occupied by the pathogen lineage on host tissue). Within-host competition effect (*c*) assumed to be determined by the most aggressive lineage (i.e., the one with the fastest lesion growth) and mixed lineage infection is assumed to have no synergistic effect on *LGR* (e.g., in wheat pathogen *Zymoseptoria tritici*, Barrett et al. 2021). Thus, the ‘competitor’ is defined as the lineage with the greatest *LGR*. Where there are multiple competing lineages for any focal lineage, the competitive effect (*c*) on the focal lineage is assumed to come from the lineage with the greatest *LGR*. The competition effect is given by:

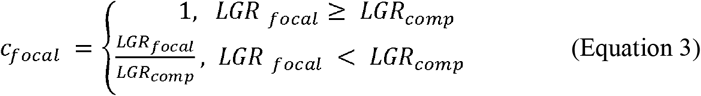

Where *LGR_focal_* and *LGR_comp_* is the *LGR* of the focal lineage and that of the competitor, respectively.

Across host species transmission (β) is defined by infection efficiency (*Y*) × number of sporangia per unit time per host (σ), where *Y* = frequency of spore contact with another host (*g*) × the infection success rate (*v*), and *σ* = *SD·LGR* (Montarry et al. 2010). Different relationships between the frequency of spore contact (*g*) and the pathogen population size (*N’*) were used, as this transmission function likely depends on the host-pathogen dynamics (see Hopkins et al. 2020). This is given by:

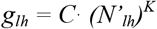

Where *C* is a scaling parameter, *K* is the strength and form of density dependence (*K* = 0 if the relationship is density independent, 0 < *K* < 1 if the relationship is non-linear, and *K* = 1 if density dependence has a strong linear relationship (modified from Hopkins et al. 2020 where the host density term is substituted by pathogen population density, and host area *A* is given a value of one since the carrying capacity is already defined through the negative density dependence term; Equation 2). This function assumes that the greater the pathogen population size, the higher the frequency of cross-host contact (*g*). Hence, *β* (cross-host transmission rate) of lineage *l* from host *h* to the other host (host *j*):

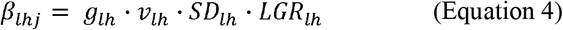

Finally, population sizes after transmission (*N*\*) just prior to the next generation (iteration) is described by:

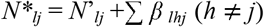

Following the stabilisation of population size, we consider the lineage proportion outputs (i.e., percentage of each lineage by the total pathogen population size within each host). The lineage proportions are of practical importance as field isolations involve random sampling of the total pathogen population, so a lineage present at very low frequency (e.g., 1 - 5%) may not be easily detected without extensive sampling.

### Modelling

Null models were first run with the following parameters to explore the behaviour of the model. The null models are: 1) all populations have equal *LGR* (of 1); 2) all populations have equal *LGR* (of 1) except a minute advantage in one lineage on one host (*LGR*_1,p_ = 1.01), or both hosts (*LGR*_1,p_ = 1.01, *LGR*_1,t_ = 1.01); and 3) same as in (2), except the advantage in that one lineage on one host is very large (*LGR*_1,p_ = 6). In these, other parameters were set to default values (Table 1) and we assume all genotypes have the same sporulation capacity (*SD* = 1). The time available for infection (*μ*), latency period (*γ*) and the carrying capacity of leaf tissue (*x*) are assumed constants and were given default values, based on those used in previous models (Table 1). These models were run with various density dependent cross-host transmission functions (*K* = 0, 0.5, and 1).

**Table 1.**
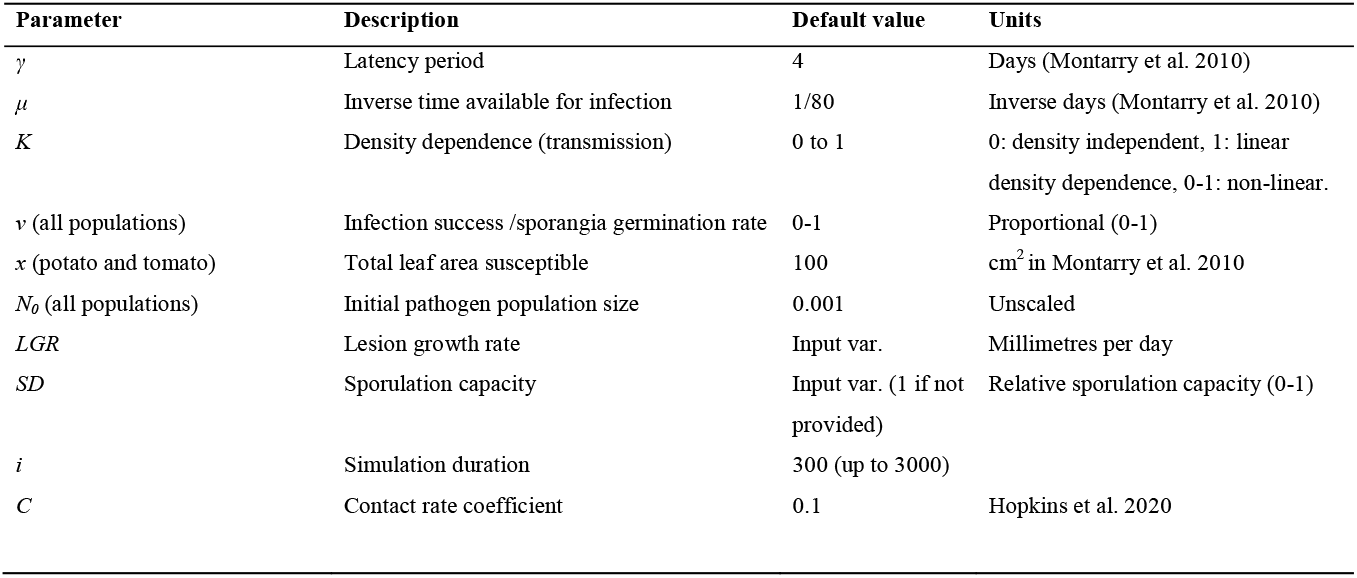
Parameters and default values used in the potato and tomato blight model. Values were changed according to data reported in studies.

To evaluate the predictive potential of the model, test data were extracted from the published pathogenicity trials on *P. infestans* lineages on both potato and tomato. Relevant trials were identified using Google Scholar using the search terms “Phytophthora infestans”, “potato”, “tomato”, and “detached leaf” (conducted early 2022). The trials tested the pathogenicity of co-occurring (and competing) lineages on both hosts under controlled laboratory conditions using detached leaf assays. GetData Graph Digitizer software (v. 2.26.0.20, available: getdata-graph-digitizer.com) was used to extract data from figures. The *LGR* (and *SD* data if reported by authors) of each lineage were used as inputs in the model (Tables 2 and 3). *SD* was given a value of 1 for all lineages if not reported by authors. Latency period and spore survival were imputed if reported by authors, otherwise default values were used (Table 1). All lesion size data were converted to lesion growth rate per day (*LGR* [mm per day] = lesion size [mm] / days post-inoculation). *LGR* were averaged to a single value for each lineage on each host if there were multiple values (e.g., data from multiple isolates). For studies presenting lesion areas, they did not report the width of lesions so length was estimated from the area using √mm^2^. Unfortunately, rigorous surveying of lineage proportions on hosts were not conducted in most cases, and the reports on the prevalence of lineages were mainly anecdotal only. Thus, output lineage proportions were compared with anecdotal observations made during the time, as reported in studies. These consists of descriptions of the relative presence of lineages on each host or observations of subsequent changes or displacements following the trial as reported in later literature. The model was written in R (ver. 4.2.2). The code is provided in the Supplementary Materials.

**Table 2.**
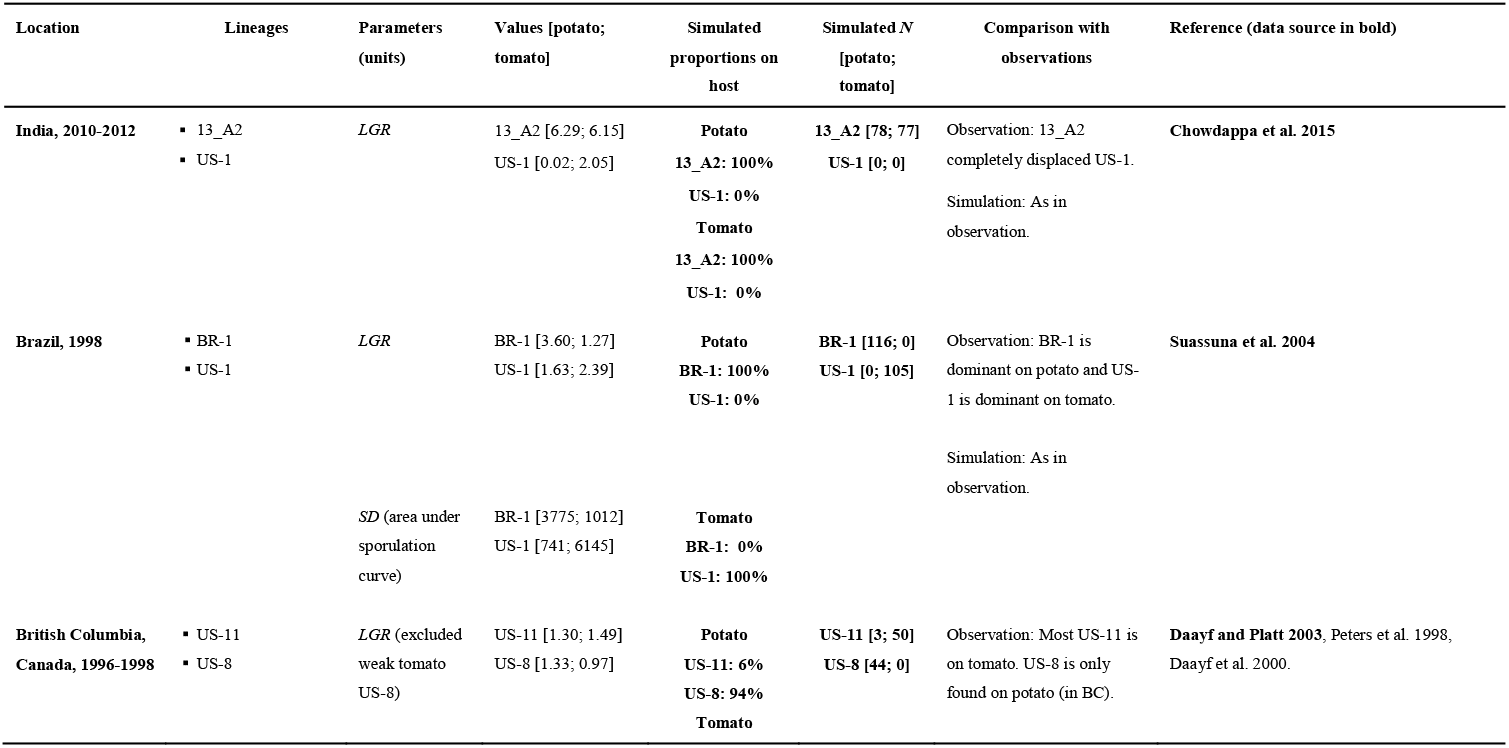
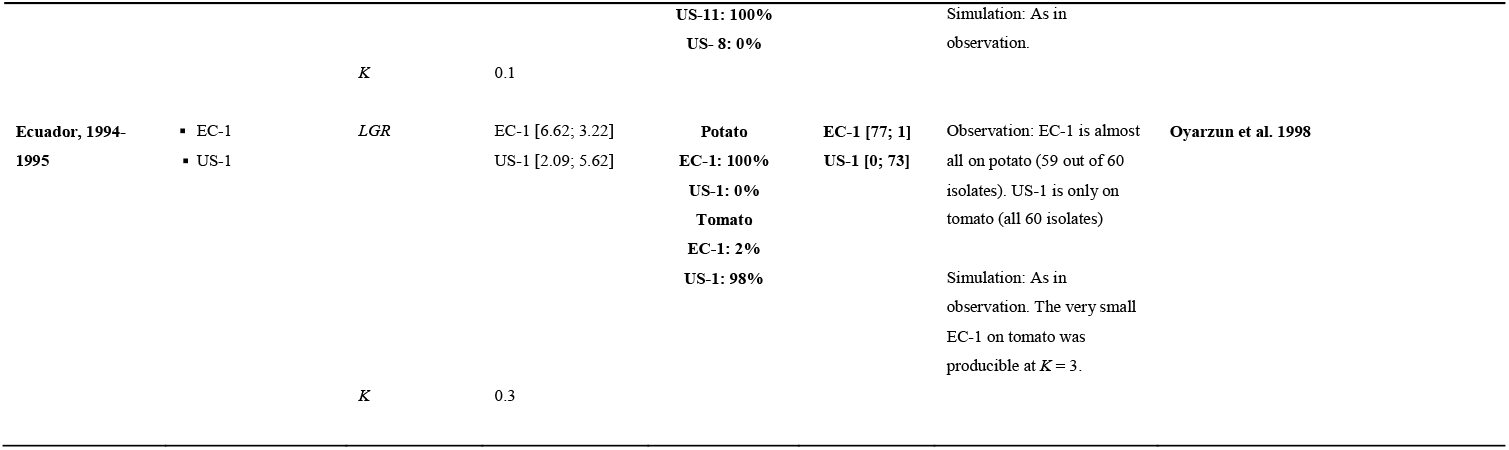
Example two-lineage simulations based on pathogen life-history trait data from studies on co-occurring *Phytophthora infestans* lineages on potato and tomato hosts. Results after 300 iterations are shown. The following constants are used unless where indicated: *γ* = 4; *μ* = 1/80, *K* = 0.5, *v* = 1, *c* = 1, *x_p_* = 100, *x_t_* = 100, *N_l,h_* = 0.001. Graphical presentations are in Fig. 1.

**Table 3.** Example simulations with three or more lineages based on pathogen trait information from studies on co-occurring *Phytophthora infestans* lineages on potato and tomato hosts. Results after 300 iterations are shown. The following constants are used unless where indicated: *γ* = 4; *μ* = 1/80, *K* = 0.5, *v* = 1, *c* = 1, *x_p_* = 100, *X_t_* = 100, *N_l,h_* = 0.001. Graphical presentations are in Fig. 2.

| Location (isolation date) | Lineages | Parameters (units) | Values [potato; tomato] | Simulated in host proportions | Simulated $N$ [potato; tomato] | Consistent with field obs.? | References (data source in bold) |
| --- | --- | --- | --- | --- | --- | --- | --- |
| Northern Algeria, 2013-2015 | 23_A1<br>13_A2<br>2_A1 | *LGR* | 23_A1 [5.02; 4.85]<br>13_A2 [4.68; 3.78]<br>2_A1 [4.62; 4.27] | **Potato**<br>**23_A1: 1%**<br>**13_A2: 99%**<br>**2_A1: 0%** | **23_A1 [1; 68]**<br>**13_A2 [95; 2]**<br>**2_A1 [0; 0]** | Observation: 23_A1 was predominantly on tomato, and 13_A2 is only on potato. 2_A1 was found mainly on tomato.<br>Simulation: Accurately predicted the complete displacement of 2_A1. However, 13_A2 should not be present at all on tomato. | **Belkhiter et al. 2019**, Beninal et al. 2021 |
|  |  | *SD* (1000 × sporangia/ml) | 23_A1 [154; 435]<br>13_A2 [489; 220]<br>2_A1 [41; 216] | **Tomato**<br>**23_A1: 97%**<br>**13_A2: 3%**<br>**2_A1: 0%** |  |  |  |
| USA and Canada, 2009-2011 | US-8<br>US-23<br>US-22 | *LGR* | US-8 [4.67; 3.04]<br>US-23 [3.76; 4.08]<br>US-22 [3.87; 4.25] | **Potato**<br>**US-8: 52%**<br>**US-23: 48%**<br>**US-22: 0%** | **US-8 [27; 0.5]**<br>**US-23 [25; 68]**<br>**US-22 [0; 0]** | Observation: US-8 is on potato only. US-22 and US-23 are mostly on tomato.<br>Simulation: Accurately predicted complete displacement of US-22 by US-23, which became prevalent on both hosts with a greater frequency on tomato. US-8 is only on potato. | **Danies et al. 2013** |
|  |  | *SD* (1000 × sporangia/ml) | US-8 [70; 19]<br>US-23 [57; 163]<br>US-22 [25; 119] | **Tomato**<br>**US-8: <1%**<br>**US-23: >99%**<br>**US-22: 0%** |  |  |  |
|  |  | *K* | 1 |  |  |  |  |
| New York State,<br>1992-1993 | ▪ US-1 | *LGR* | US-1 [4.99; 1.86] | **Potato** | **US-1 [0 ; 0]** | Observation: US-1 and US-8 are potato specific. US-7 emerged in 1992 and became completely dominant in 1993. | **Legard et al. 1995** |
|  | ▪ US-7 |  | US-8 [5.14 ; 1.80] | **US-1: 0%** | **US-8 [0 ; 0]** |  |  |
|  | ▪ US-8 |  | US-7 [5.20 ; 4.27] | **US-8: 0%**<br>**US-7: 100%** | **US-7 [44 ; 134]** |  |  |
|  |  |  |  |  |  | Simulation: Accurately predicted the complete dominance of US-7 (*K* > 0.7). At *K* = 0.5, potato is shared among US-8 (72%) and US-7 (28%). |  |
|  |  | *SD* (1000 × sporangia/ml) | US-1 [367; 10]<br>US-8 [458; 6]<br>US-7 [346 ; 198] | **Tomato**<br>**US-1: 0%**<br>**US-8: 0%**<br>**US-7: 100%** |  |  |  |
|  |  | *K* | 0.8 |  |  |  |  |

### Analytical solutions for displacement and coexistence

The conditions for which a lineage is dominant or no lineage is dominant on all hosts is described analytically. We introduce some definitions about stability of steady states. Consider an iteration equation:

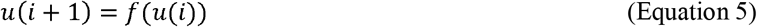

Where *u* = (*u*_1_, *u*_2_,…, *u_N_*) and *f*: ***R**^n^* → ***R**^N^*. We call *u*\* is a *steady state* of (5) if *u*\* = *f*(*u*\*). *u*\* is *trivial* if *u*\* = (0,0,…,0) and *u*\* is *semi-trivial* if *u*\* is not trivial and there is an 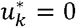 for some *k*. Moreover, a steady state *u*\* is a *coexistence* if 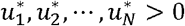. A steady state *u*\* of (5) is *locally stable* if there is an *δ* > 0 such that if |*u*(0) – *u*\* | < *δ*, then *u*(*i*) → *u*\* as *i* → ∞.

For *m* lineages and *n* hosts (*m, n* ≥ 2) where 1 ≤ *l* ≤ *m* and 1 ≤ *h* ≤ *n*,

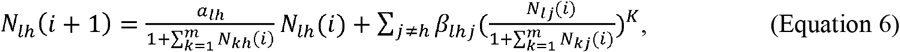

Where *a_lh_, β_lhj_* > 0 for all 1 ≤ *l* ≤ *m*, 1 ≤ *h* ≤ *n* and *j* ∈ {1,2,…, *n*} – {*h*}.

#### Theorem 1

For 0 < *K* < 1,

a. For any initial data *N_lh_*(0) ≥ 0, we have *N_lh_*(*i*) ≤ *w_lh_* for all 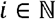, where

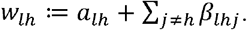
b. If 0 < *N_lh_*(0) ≤ *w_lh_* for all 1 ≤ *I* ≤ *m* and 1 ≤ *h* ≤ *n*, then

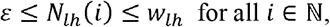

where 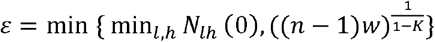 for 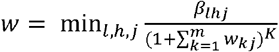.
c. All trivial and semi-trivial steady states of (6) are not locally stable.
d. There is a coexistence in (6).

Proof: For (a), it follows from

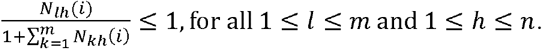

Next, we prove (b) by induction. Indeed, from the definition of *ε*, we have *ε* ≤ *N_lh_*(0) ≤ *w_lh_* and

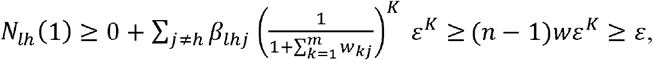

where the last inequality holds by using the definition of *ε*. Hence, by (a), we obtain

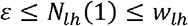

Continuing this process, we get *ε* ≤ *N_lh_*(2) ≤ *w_lh_*, and so on. We complete (b).

For (c), we can choose any initial pathogen population size *N_lh_*(0) > 0, being very closed to a trivial or semi-trivial steady state. From (b), we have *N_lh_* (*i*) ≥ *ε* for all 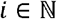, which implies 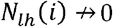 as *i* → ∞. Hence a trivial and all semi-trivial steady states are not locally stable.

For (d), we identify *u* = (*u*_1_, *u*_2_,…, *u_mn_*) = (*N*_11_, *N*_12_,…,*N*_1*n*_, *N*_21_, *N*_22_,…, *N_mn_*). Consider a set 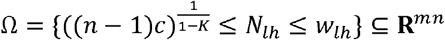

Using (b), we can define a function *f*: Ω → Ω by

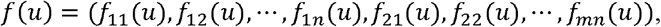

where

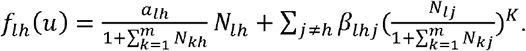

Note that *f* is continuous from *Ω* to *Ω*. Since *Ω* is compact, we can apply the Brouwer fixed point theorem to insure the existence *u*\* ∈ *Ω* such that *u*\* = *f*(*u*\*). Therefore, *u*\* is a coexistence of (6).

Applying Theorem 1 to 2 lineages and 2 hosts (*m* = 2 and *n* = 2). For 1 ≤ *l* ≤ 2 and 1 ≤ *h* ≤ 2, we have:

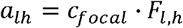

and

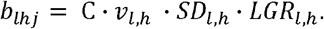

#### Definition 1

We consider the *h*-th lineage is dominant in the *h*-th host if:

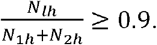

The result below gives a sufficient condition to ensure that no lineage is dominant in all hosts.

#### Corollary 1

Set

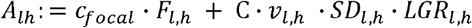

and

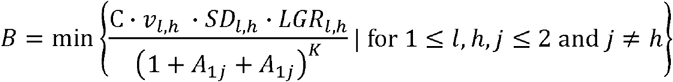

If

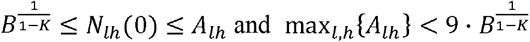

then no lineage dominates in all hosts for all stages.

*Proof*

By the assumption and Theorem 1 (b), we obtain:

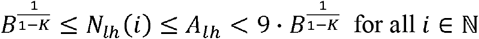

and hence,

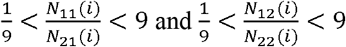

Consequently, for all 1 ≤ *l* ≤ 2 and 1 ≤ *h* ≤ 2 and all 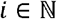

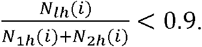

## Results

The results from the null models show that the lineage proportions were distributed evenly (i.e., 1/*l*) on each host in the case where each lineage have the same pathogenicity traits (are equally fit). Giving one lineage a minute advantage (e.g., a greater *LGR* of only 0.01 mm/day) on either or both hosts showed that the lineages with lower fitness were not completely displaced unless there is strongly density dependent cross-host transmission (e.g., *K* > 0.75). Giving one lineage a very large advantage on one host (e.g., lineage 1 on potato with the advantage: *LGR*_1,*p*_ = 6, all others = 1) resulted in complete displacement of other lineages for most transmission functions (e.g. *K* > 0.35). For lower or no density dependent transmission, displacement occurs within the host with that large *LGR* difference only (i.e., potato in that example). Simulations showed that the fitness estimates from empirical *LGR* and *SD* can replicate the historical observed genotypic compositions except in a few cases. Fitting other pathogenesis parameters such as latency period (where the data was available) did not have large influences on the final lineage proportions so model results with default values were presented. Data on the field lineage frequencies was unavailable to rigorously assess the accuracy of fitness estimates, so the model is necessarily heuristic in seeking to simulate past epidemic situations. Nonetheless, model outputs are generally consistent with the observed historical epidemiology.

## Simulations involving two lineages

The results from the two-lineage models are summarised in Table 2 and Fig. 1. Oyarzun et al. (1998) described an interaction between US-1 and a newer invader EC-1, which infect tomato and potato respectively. A survey conducted from 1993 to 1996 found that all isolates of US-1 were from tomato (60 out of 60), and all but one of EC-1 were from potato (59 out of 60) with one on tomato (1 of 60 = 1.7%). Simulations reproduced the expected strong host specificities of EC-1 and US-1 using the *LGR* data presented. At *K* = 0.5, EC-1 formed 99% and 4%of the population on potato and tomato respectively, while US-1 formed 1 vs. 96% on potato and tomato respectively. At *K* = 0, both lineages are completely dominant on their hosts. At *K* = 1, lineages coexist on both hosts (41% EC-1 and 59% US-1 on tomato), but EC-1 is very dominant on potato (96% EC-1 and 4% US-1). *K* = 0.3 reproduces the approximate lineage frequencies reported in the study (Table 2).

**Fig. 1.**
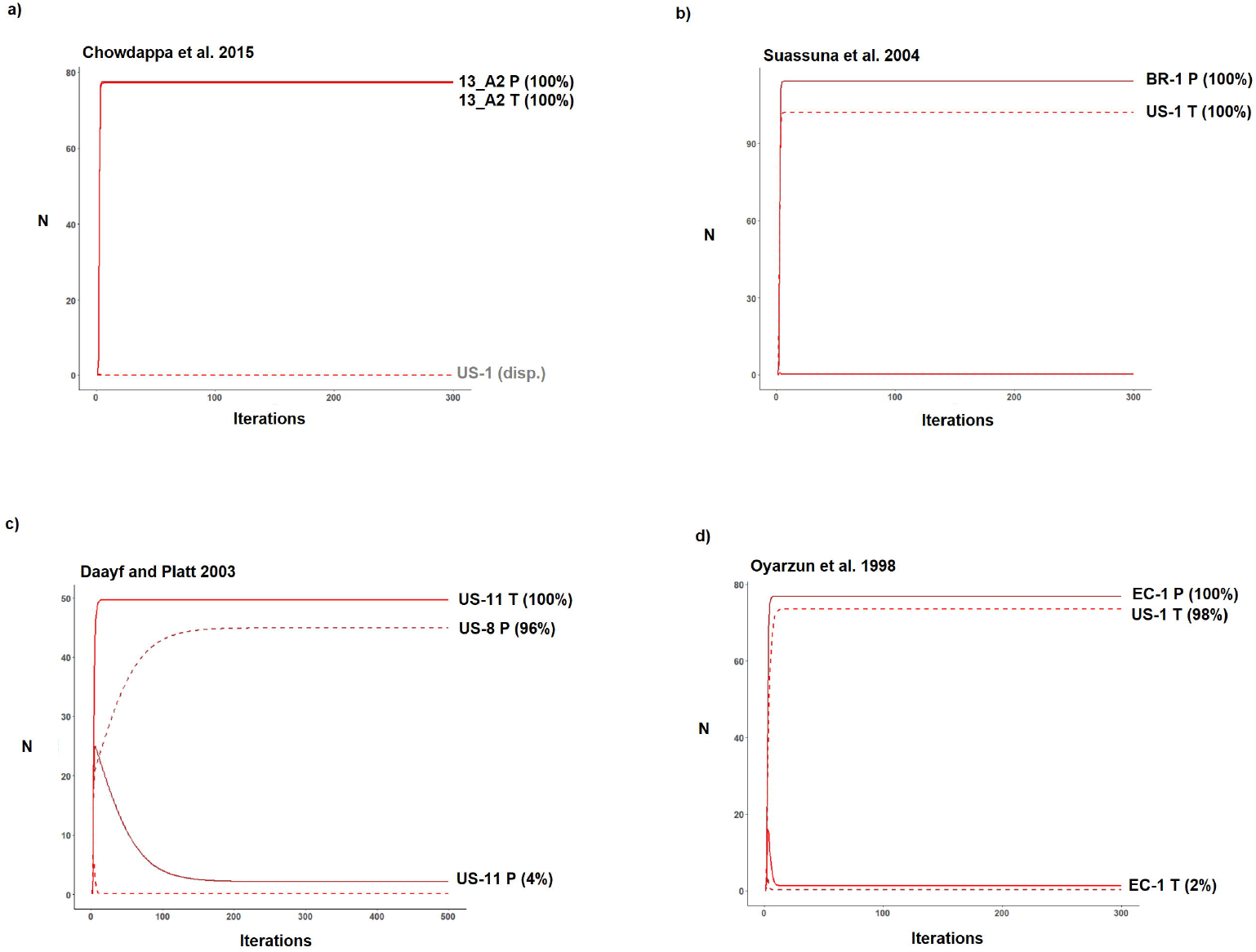
Two-lineage simulations on potato and tomato populations of *Phytophthora infestans* on potato and tomato (*N* is the pathogen population size). Simulations used empirical data from the published data on pathogenesis parameters of lineages on the two hosts. Red and brown lines indicate pathogen populations on tomato and potato respectively. Solid lines and dashed lines denote lineages as labelled on the lines. Displaced lineages are shown in grey text.

Daayf and Platt (2003) tested isolates of US-8 and US-11 which co-occurred mostly in British Columbia (BC), Canada, in 1996-1998. Detailed information on the proportions of lineages on each host at sites during this period was not available. During 1996-1999, potato blight was much more prevalent than tomato blight, the latter only impacting a few Canadian provinces (mostly BC) where tomatoes were mainly grown in home gardens (Peter et al. 1998; Daayf et al. 2000; Daayf and Platt 2003). All tomato isolates were of the A1 mating type (US-11) except on Prince Edward Island (PEI) and Nova Scotia which were US-8 (A2). Isolates from different provinces were pooled and tested in the study. Sporulation was not reported and the simulations were run only with *LGR*. In 1996, US-11 was reported on both hosts in British Columbia and was only found on tomato in Ontario (Daayf et al. 2000). The US-11 seemed to be displaced by US-8 by 1998, but there were no details on the host of origin of isolates and there was low sampling (Daayf et al. 2000). Overall, it was expected that the results should show US-11 dominant on tomato with a small potato population, and US-8 should be only on potato (US-8 tomato populations were only found in Prince Edward Island and Nova Scotia). In the simulations, this was replicated at *K* = 0. Under strong density dependence (*K* = 1), US-8 is completely displaced by US-11 on both hosts. Then, it would appear that demographic barriers were limiting the spread of the generalist US-11 (such as the relatively limited availability of tomato in home gardens), or that the pooled data of isolates from multiple provinces (including BC) were not representative of populations at BC so the interaction between those lineages could not be modelled. In any case, there was a lack of sporulation data for constructing fitness.

From Brazil, Suassuna et al. (2004) reported that BR-1 was only found on potato and US-1 was only on tomato, where BR-1 (A2 mating type) is a new invader emerging in the mid-80s. Potato was grown at higher altitudes (> 800m) and tomato crops was more widespread than potato. The isolates tested were collected in 1998. The finding that BR-1 was specific to potato and US-1 was specific to tomato was replicated under all density dependence scenarios (all *K*).

Legard et al. (1995) described two major migration events in North America (i.e., US-1 followed by US-6 in the late 1970s). US-6 was mainly associated with tomatoes and occasionally potatoes and was metalaxyl resistant, while US-1 was almost entirely associated with potato and was metalaxyl sensitive (Platt 1999). In this region, US-6 was the most widespread genotype during this period (Goodwin et al. 1994; but not in New York, Goodwin et al. 1998). However, populations (fields) only consisted of a single clonal lineage at each location (Goodwin et al. 1994) so they might not have co-occurred at the same sites. A hypothetical interaction was simulated between the generalist US-6 and the older potato-specific US-1 using data from lineages isolated from 1987 to 1991 (Table S1). US-6 was introduced in the late 1970s and was most common on tomato in Mexico, and may have caused outbreaks on tomato in California in 1979. Appropriate data on the population frequencies of lineages at the time was not available (Goodwin et al. 1994) but it may be reasonably assumed that US-1 was not competitively displaced by US-6. US-6 is completely dominant on tomato (100%) but both coexist on potato at *K* = 0.5. However, US-6 completely displaces US-1 on both hosts at *K* = 1.

Without density dependent transmission (*K* = 0), US-6 is completely dominant on tomato, and US-1 is very dominant on potato (97%) with a low frequency of US-6 (3%). Thus, in all cases US-1 was completely excluded from tomato by US-6.

## Simulations with three or four lineages

The results from the lineage models with three or four lineages are presented in Table 3 and Fig. 2. Danies et al. (2013) studied US-22, US-23, US-24, and US-8 strains isolated during 2009 to 2011 in locations across North America. US-22 spread quickly in 2009, just a few years after emergence. Subsequently US-23 and US-24 emerged within that same year (Saville and Ristaino 2019). US-24 was limited to colder regions such as North Dakota and Wisconsin (Hu et al. 2012). The US-23 displaced the remaining US-22 on tomato in 2012, while US-8 remained persistent on potato at ~20% of isolates across years (as of 2016, Saville and Ristaino 2019). The three-lineage simulations involving US-8, US-22, and US-23 showed that US-22 was completely displaced (*N* ≈ 0) under all parameters of *K*. US-23 has low frequency on potato at *K* = 0.5 (3%), with US-8 very dominant (97%). At *K* = 0, US-23 and US-8 are segregated in tomato and potato respectively. At *K* = 1, the potato population consist of mixed US-23 (48%) and US-8 (52%) while US-23 is close to completely dominant on tomato (~100%). In a hypothetical interaction involving all four lineages (adding US-24), simulations showed a similar result, with US-24 also being displaced under all *K*. These results are consistent with historical observations that US-22 quickly became extinct and US-24 was restricted to some northern areas where US-23 was not present. However, the simulated proportion of US-8 should have been lower than that of US-23 (i.e., 2011-2016, see Fig. 1b and 1c of Saville and Ristaino 2019). The decline of US-8 relative to US-23 was attributed to either reduced mefenoxam usage (to which the former was resistant but not the latter) or increasing pathogenic fitness in US-23 in the years following that pathogenicity trial (Saville and Ristaino 2019), which could explain the inflated prevalence of US-8 over US-23 in the model. Nonetheless, the pathogenesis data from 2009 accurately predicted the displacement of US-22 in the subsequent one or two years (2010-2011). Unfortunately, the pathogenicity of US-11 (which was also extant at relatively low frequency) was not tested.

**Fig. 2.**
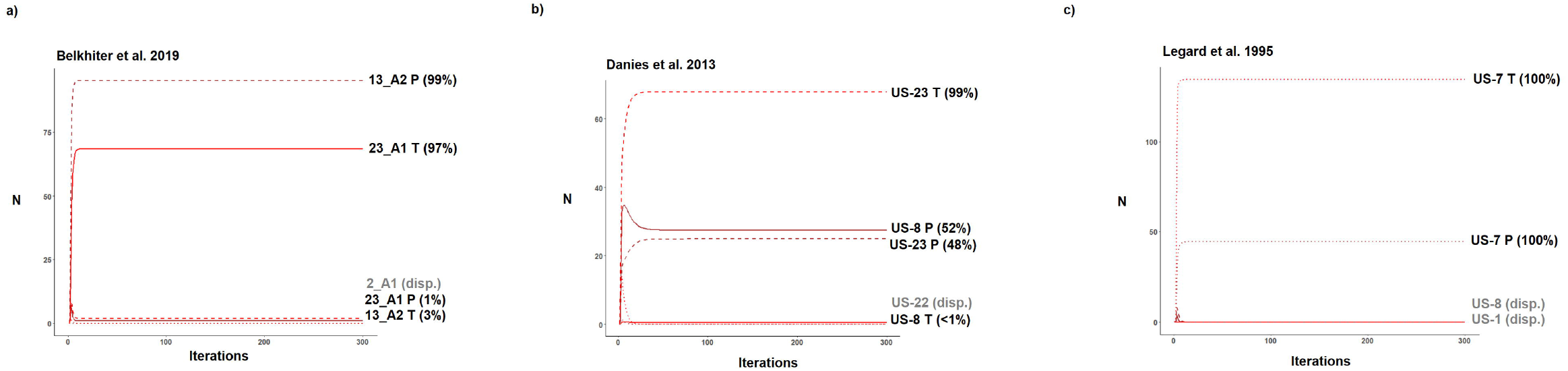
Three-lineage simulations on potato and tomato populations of *Phytophthora infestans* on potato and tomato (*N* is the pathogen population size). Simulations used empirical data from the published data on pathogenesis parameters of lineages on the two hosts. Red and brown lines indicate pathogen populations on tomato and potato respectively. Displaced lineages are shown in grey text.

Belkhiter et al. (2009) tested isolates of three predominant lineages in northern Algeria (2_A1, 13_A2, and 23_A1) during 2013 to 2015. 2_A1 was the oldest lineage and was eventually displaced by 13_A2. In the previous period of 2008 to 2012, 2_A1 was the most common and mainly infects potato. Following 2013, 13_A2 became the most dominant but 2_A1 was not displaced (Rekad et al. 2017). 23_A1 was first detected in 2013 and is associated with both hosts (but more with tomato) at low frequency. It was the least commonly isolated among the three and was only found in some locations around Mostaganem. The simulations were based on isolates mostly recovered during 2014 to 2015. They show that at *K* = 1, the population on potato is mainly 13_A2 (96%) and partially 23_A1 (4%); and tomato population is a mixture of the two (64% 23_A1 and 36% 13_A2). At *K* = 0, 13_A2 and 23_A1 are completely dominant on potato and tomato, respectively. At *K* =0.5, 13_A2 and 23_A1 are 99% and 1% on potato and 97% and 3% on tomato, respectively. 2_A1 is completely displaced in all scenarios. In surveys 13_A2 was not observed on tomato, but the simulations inaccurately suggested that it generally can sustain a population on tomato. In the short-term, the complete displacement of 2_A1 in the output appeared to be inconsistent with the observed proportions because 2_A1 was still very common in 2014 (Rekad et al. 2017). However, 2_A1 subsequently declined and was not recovered later in 2016 (Beninal et al. 2021) consistent with the model output.

Lastly, the test data (isolates from 1987 to 1991) of Legard et al. (1995) was used to simulate the situation in New York State during 1992 to1993. The proportions in 1992 for US-7, US-8 and US-1 were 55 isolates, 43 isolates and 4 isolates, respectively. In 1993, only US-7 was found in the state (Goodwin et al. 1995). The simulations demonstrated that at *K* = 0, the potato population consisted of US-8 (98%) and US-7 (2%), but US-7 was completely dominant on tomato (100%). At *K* = 0.5, the potato population was US-8 (72%) and US-7 (28%), and US-7 was also completely dominant on tomato (100%). Strong density dependence at *K* = 1 (or > 0.75) produced results that reflected subsequent field observations, where US-7 was completely dominant on both hosts and all other lineages displaced. Interestingly, US-1 was completely displaced in all the scenarios. This result on the displacement of US-1 and US-8 by US-7, along with other results on the displacement of US-22 from Danies et al. (2013) and 2_A1 from Belkhiter et al. (2019) indicate that fitness calculated from pathogenesis trial data could be used to project future population changes, even when the displaced lineages were common during the time of isolate collection. Another potential utility is to predict the outcome of interactions among lineages in hypothetical interactions (Table S1). Whether a generalist or specialist genotype is displaced in the two-lineage model (where all else is equal except for *LGR*) is presented in Figure 3.

**Fig. 3.**
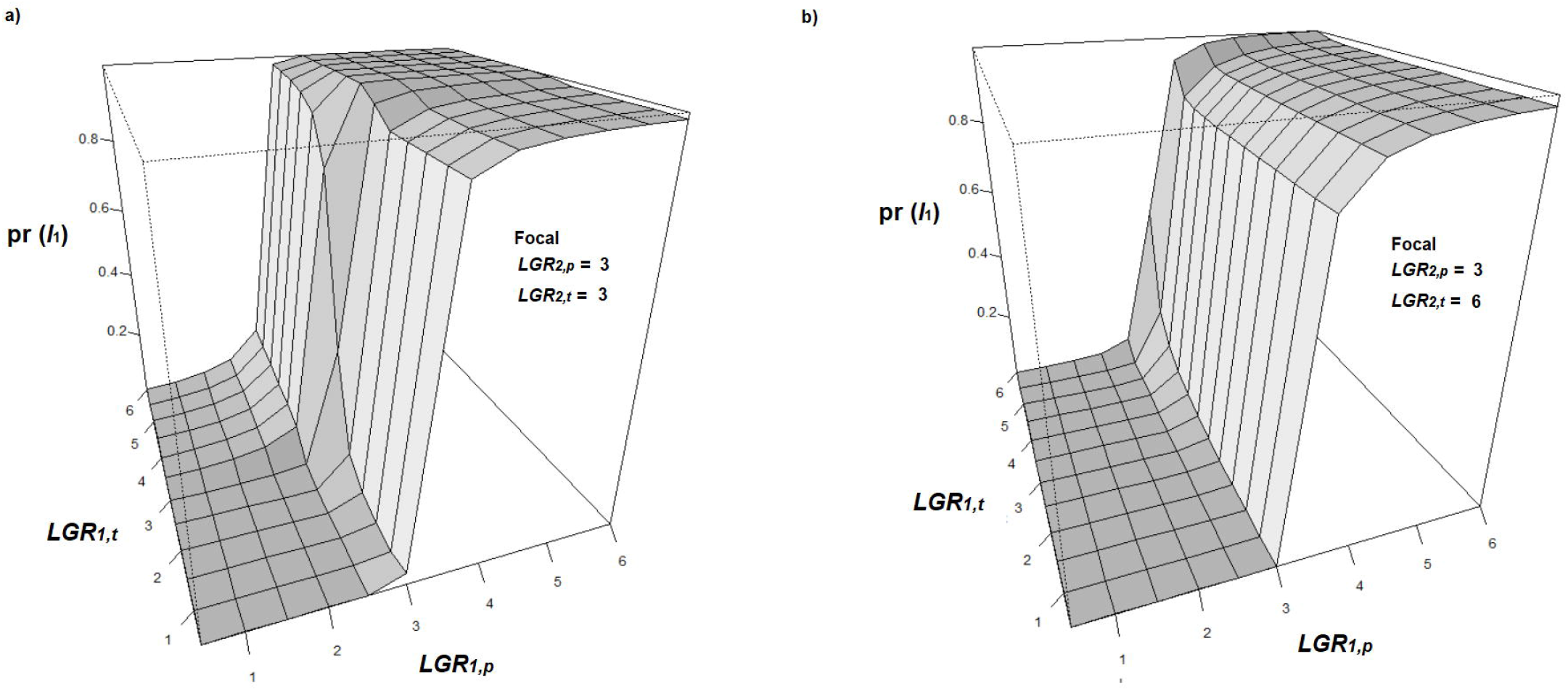
Three-dimensional graphs of the simulated final frequency of Lineage 1 on potato in the presence of competing Lineage 2: pr (*l*_1*p*_) from the model involving two hosts (i.e. potato and tomato). Results are following stabilization after 300 iterations. Sporulation capacity (*SD*) of both lineages is assumed equal and all other input parameters are as in Table 1. The results from Lineage 1 with various lesion growth rates (*LGR*) on potato (*p*) and tomato (*t*) are presented. The proportion of Lineage 2 (*l*_2_) is [1 – pr (*l*_1_)]. In panel a), the competitor is a generalist with *LGR* = 3 on both hosts. In panel b), the competitor is a specialist with *LGR* = 6 on tomato and *LGR* = 3 on potato.

## Discussion

The goal of this work is to develop a simulation model for clonal pathogen lineage frequencies on multiple host species based on pathogenesis traits, providing a tool for predicting fluxes in populations of important plant pathogens such as *P. infestans*. The genotypic fitness on each host was calculated from the pathogenesis parameters derived from empirical tests of *P. infestans* on potato and tomato, in particular lesion growth rate and sporulation capacity (Montarry et al. 2010). Using a demographic model, fitness was used to project population changes. With this model, the interaction between genotypic fitness and pathogen population biology can also be assessed. As such, the model has practical and theoretical applications alongside knowledge on host specificities and fungicide sensitivities of extant lineages at a site (Fry et al. 2015). The current model differs from the previous model (Yuen 2013) on lineage displacements, in that its construction is a demographic simulation model based on pre-existing models on pathogen fitness and population demography. Furthermore, this model focuses on two major hosts (i.e., potato and tomato) instead of one. Lineage persistence is deterministic based on relative fitness, so a pre-existing dominant lineage would not have any effect on the final frequencies of a new invader with small population size (i.e., output is not affected by the starting population size). This model behavior is reasonable as previous experiments suggest that competition among *P. infestans* strains is extreme, such that differences in initial starting inoculum size may not influence the final outcome of competitive interactions (Young et al. 2009). For instance, a competition experiment found that in the absence of fungicides, competitive exclusion by a more aggressive strain (i.e., with faster lesion growth) can occur even when that strain started at 1% frequency (Kadish and Cohen 1988).

In general, the model demonstrates that complete displacements occur when the fitness difference between lineages is sufficiently large within both hosts. The relationship between pathogen population size and transmission rate is important. A systematic review of host-parasite systems (mostly in vector or sexual contact diseases) found that most empirical studies report non-linearity in this relationship (Hopkins et al. 2020), so *K* = 0 or 1 are considered uncommon. The assumption that lesion growth rate is the determining factor for the competitive effect is reasonable for *P. infestans* because there is ample evidence that lineages with fast lesion growth displace slower ones (e.g., Miller et al. 1998; Gisi et al. 2011; Njoroge et al. 2018; but see Mariette et al. 2016). However, this may be inappropriate for other pathogen species that have different resource consumption strategy, such as those with strains that infect plants at different ages so high lesion growth rate may not always the most favorable (Dutt et al. 2021).

The simulation results are typically in line with field observations of the population make-up of *P. infestans* lineages on potato and tomato. Interestingly, the model retrospectively predicted the displacements of US-22 in Danies et al. (2013) and 2_A1 in Belkhiter et al. (2019) before their occurrence a year or few years prior. Likewise, the complete dominance of US-7 on both hosts in New York in 1993 (displacing co-occurring US-1 and US-8, Goodwin et al. 1995) was reproduced using the data from isolates recovered from the field at the time (Legard et al. 1995). This suggests that the empirical pathogenesis data can have a high degree of accuracy in estimating genotypic fitness through the well-established mathematical definition of pathogen fitness (Montarry et al. 2010). Notably, the displaced lineages (e.g., US-22 and 2_A1) were very prevalent at the time of isolation, but the model accurately predicted they would be completely displaced by competing lineages. The model showed some promise in predicting small lineage frequencies from the fitness estimations (i.e., 1-5% on host), such as the rare EC-1 on tomato in Ecuador at the time (Oyazun et al. 1998). These together highlight the potential validity of the pathogenicity parameters in estimating fitness.

Sporulation capacity is an important trait in fitness and in the competitive interaction among strains (Kadish and Cohen 1988; Montarry et al. 2010). Unfortunately, this was not always tested in trials. There are several factors that were unaccounted for in the model that can affect accuracy. Barriers to dispersal and host density asymmetry are beyond the scope of the model, but can have important effects on dispersal and transmission. The current model assumes a high density of hosts so the rate of cross-host infection increases with the amount of disease in the field (e.g., Biere and Honders 1998), which may be appropriate for adjacent potato and tomato farms. This may not be suitable in cases where transmission rate is more host-density dependent such as natural ecosystems (Antonovics 2017). An infrequent lineage on an uncommon host might also be more difficult to recover if sampling is not intensive enough. A direct spatial design or a model that explicitly considers host asymmetry is likely necessary for some regions. For instance, the model incorrectly indicated a low frequency of Algeria 13_A2 on tomato (Table 3), but 13_A2 was only found on potato after extensive sampling (note that 13_A2 is aggressive on both hosts in India; Chowdappa et al. 2015). In Algeria, the area of potato production is much greater than tomato (156k ha for potato versus 22k ha for tomato). Potato is grown in three seasons while tomato is grown year-round in plastic houses and fields (Belkhiter et al. 2019). If many tomato plants are inside plastic houses, then there may be significant dispersal barriers for cross-host transmission from potato to tomato. As such the host availability of tomato may be much lower than was assumed in the model. The general applicability of the current model can be tested in field trials in future work.

There are challenges in accounting for host density asymmetry, for which the model does not explicitly consider. For instance, changing the host tissue available for infection *x_h_* (following the ratio 22 for tomato and 156 for potato) and season length *μ_h_* (60 for tomato and 80 for potato) produced outputs similar to that of the original model, because the effect of changing *x* on fitness is minimal unless *x* is very small (< 10). However, lowering *x_t_* to 5 suggests 13_A2 should have high frequency on tomato (> 20%) in the model, which does not reflect the field situation at that location. Modelling of the epidemic biology in Algeria needs to take into account multiple plantings in a year (main and late season) because 23_A1 is infectious during the late season only (Beninal et al. 2021), so their competitive effect may not be as strong as indicated in the model. Similarly, tomato plants were grown in home gardens and not in fields in Canada (Daayf and Platt 2003), so tomato might have much lower density than potato. Lowering *x_t_* (e.g., to 5) while keeping *K* = 0.1 showed a small population of US-8 on tomato (which does not reflect the situation at British Columbia), and on potato the output was similar to that of the original model. At *K* = 0.5 there is coexistence between US-11 and US-8 on tomato. Interestingly, tomato US-8 was found at relatively high frequency on PEI (Daayf and Platt 2003), from which some of the US-8 isolates were sourced from and pooled with other states. The absence of US-11 on PEI might indicate that the lineage was never introduced there or there was insufficient tomato density. There was no sporulation data available which likely affect the overall precision of the results. Overall, it is important that the isolates recovered should be representative of the focus study area. Adjacent areas may have contrasting population compositions on potato and tomato which may stem from differential fitness among populations from different geographical locations (i.e., variations within lineages; Mariette et al. 2016).

Other caveats are that replication in trials should be sufficient to reduce variability in the trait means, so caution should be given where the trait means of lineages are close. There is possibility that relative differences in pathogenesis traits among lineages may be incorrect due to data variance, which can affect the outcome of simulations. This includes variation in aggressiveness of lineages between seasons. The current model does not consider temporal scales explicitly and only has the assumption that lineages are co-occurring and competing at some given location. Displacements often occur very quickly in *P. infestans* (between two seasons), but the model does not consider the length of time required for displacement. It assumes that displacement occurs quickly on an unscaled time frame and does not focus on short-term fluxes in initial iterations. Genotype (or lineage) × cultivar interaction effects might affect the outcome of competition (e.g., Young et al. 2009), so the proportions of cultivars with varying disease resistance in the field could determine the frequency of lineages (Stellingwerf et al. 2018). There may be cases where the strain with the highest lesion growth rate and sporulation density may not be the most successful, possibly due those temporal changes in host availability or trade-offs between vertical and horizontal transmissions (e.g., French 13_A2 on potato, Mariette et al. 2016). However, both potato and tomato hosts should be considered when modelling fitness. This issue can be addressed in extensions of this model.

In future models on lineage displacements, larger-scaled spatial simulation models can take into account short- and long-range dispersal, host infection dynamics and environmental parameters. Evolutionary or temporal changes in pathogenicity and the effects of fungicide on fitness are particularly emphasized to improve the accuracy of models (Saville and Ristaino 2019). Other exciting areas for development include the seasonality of cropping in areas that permit more than one potato planting per year. Coexistence could be solely explained by one strain performing better in a growing season than the other strain and the other way around (modelled in Mailleret et al. 2011). While simulation modelling is useful in epidemiology, like in all other models the factors are grossly simplified (such as competitive interactions) compared to reality (Yuen 2012). Empirical tests on the relationship between traits and multi-seasonal fitness are needed for insights on the results of competitive interaction between pathogen lineages.

## Acknowledgements

Thank you to Stephen Bonser for his guidance on the demographic modelling of populations.

