## Supplementary material for "Modelling the displacement and coexistence of clonal lineages of *Phytophthora infestans* through revisiting past outbreaks": Table S1

**Supplementary Materials**

**Table S1**

Simulations based on hypothetical interactions involving lineages that did not co-occur at the same geographical location.

| Location | Lineages (field specificity) | Parameters (units) | Values [potato; tomato] | Simulated in host proportions | Simulated *N*  [potato; tomato] | References |
| --- | --- | --- | --- | --- | --- | --- |
| USA and Canada 1987-1991 | - US-1 - US-6   Did not co-occur in NA (Goodwin et al. 1994). | *LGR* | US-1 [4.99; 1.86] US-6 [4.95; 4.37] | **Potato  US-1: 60% US-6: 40%** | **US-1 [45; 0] US-6 [30; 138]** | Legard et al. 1995 |
|  |  | *SD* (1000x sporangia per lesion) | US-1 [367; 10]  US-6 [318; 225] | **Tomato**  **US-1: 0%**  **US-6: 100%** |  |  |
| USA and Canada, 2009-2011 | - US-8 - US-23 - US-22 - US-24   US-24 in North Dakota only  US-22 and US-24 displaced by US-23 during 2019. | *LGR* | US-8 [4.67; 3.04]  US-23 [3.76; 4.08]  US-22 [3.87; 4.25]  US-24 [3.21; 1.96] | **Potato**  **US-8: 56%**  **US-23: 44%**  **US-22: 0%**  **US-24: 0%** | **US-8 [39; 0]**  **US-23 [12; 71]**  **US-22 [3; 3]**  **US-24 [0; 0]** | Danies et al. 2013 |
|  |  | *SD* (1000 × sporangia/ml) | US-8 [70; 19]  US-23 [57; 163]  US-22 [25; 119]  US-24 [30; 3] | **Tomato**  **US-8: 0%**  **US-23: 100%**  **US-22: 0%**  **US-24: 0%** |  |  |
|  |  | *v* (direct sporangia germination rate, 15ºC) | US-8 [0.48]  US-23 [0.75]  US-22 [0.45]  US-24 [0.60] |  |  |  |
|  |  | *K* | 0.9 |  |  |  |
| USA and Canada 1987-1991 | - US-1 - US-6 - US-7 - US-8   The lineages did not all co-occur at one location. | *LGR* | US-1 [4.99; 1.86]  US-6 [4.95; 4.37]  US-8 [5.14 ; 1.80]  US-7 [5.20 ; 4.27] | **Potato**  **US-1: 0%**  **US-6: 2%**  **US-8: 0%**  **US-7: 98%** | **US-1 [0 ; 0]**  **US-6 [1 ; 67]**  **US-8 [0 ; 0]**  **US-7 [64 ; 6]** | Legard et al. 1995 |
|  |  | *SD* (1000 × sporangia/ml) | US-1 [367; 10]  US-6 [318; 225]  US-8 [458; 6]  US-7 [346 ; 198] | **Tomato**  **US-1: 0%**  **US-6: 92%**  **US-8: 0%**  **US-7: 8%** |  |  |
